# Controlling the Rate of GWAS False Discoveries

**DOI:** 10.1101/058230

**Authors:** Damian Brzyski, Christine B. Peterson, Piotr Sobczyk, Emmanuel J. Candés, Malgorzata Bogdan, Chiara Sabatti

## Abstract

With the rise of both the number and the complexity of traits of interest, control of the false discovery rate (FDR) in genetic association studies has become an increasingly appealing and accepted target for multiple comparison adjustment. While a number of robust FDR controlling strategies exist, the nature of this error rate is intimately tied to the precise way in which discoveries are counted, and the performance of FDR controlling procedures is satisfactory only if there is a one-to-one correspondence between what scientists describe as unique discoveries and the number of rejected hypotheses. The presence of linkage disequilibrium between markers in genome-wide association studies (GWAS) often leads researchers to consider the signal associated to multiple neighboring SNPs as indicating the existence of a single genomic locus with possible influence on the phenotype. This a posteriori aggregation of rejected hypotheses results in inflation of the relevant FDR. We propose a novel approach to FDR control that is based on pre-screening to identify the level of resolution of distinct hypotheses. We show how FDR controlling strategies can be adapted to account for this initial selection both with theoretical results and simulations that mimic the dependence structure to be expected in GWAS. We demonstrate that our approach is versatile and useful when the data are analyzed using both tests based on single marker and multivariate regression. We provide an R package that allows practitioners to apply our procedure on standard GWAS format data, and illustrate its performance on lipid traits in the NFBC66 cohort study.

## Introduction

In the last decade, genome-wide association studies (GWAS) have been the preferential tool to investigate the genetic basis of complex diseases and traits, leading to the identification of an appreciable number of loci [1]. Soon after the first wave of studies, a pattern emerged: there exists a sizable discrepancy between, on the one hand, the number of loci that are declared significantly associated and the proportion of phenotypic variance they explain [2] and, on the other hand, the amount of information that the entire collection of genotyped single nucleotide polymorphisms (SNPs) appears to contain about the trait [3, 4]. In order to increase the number of loci discovered (and their explanatory power), substantial efforts have been made to obtain larger sample size by genotyping large cohorts [5, 6] and by relying on meta-analysis. However, the gap remains, although not as large as in the original reports. This parallels, in part, the discrepancy between the multivariate model that is used to define complex traits and the univariate approach to the discovery of associated SNPs which is standard practice, as underscored, for example, in [7–9].

Two approaches to bridge the gap emerge quite naturally: (a) an attempt to evaluate the role of genetic variants in the context of multivariate models, more closely matching the underlying biology, and (b) relaxing the very stringent significance criteria adopted by GWAS to control the false discovery rate (FDR) [10] rather than the family-wise error rate (FWER)—a strategy that has been shown attractive when prediction is considered as an end goal together with model selection [11]. Both strategies have been pursued, but have encountered a mix of success and challenges.

Multivariate models for the analysis of GWAS data have been proposed as early as 2008 [12, 13]: examining the distribution of their residuals, it is clear that they provide a more appropriate model for complex traits. However, their use to discover relevant genetic loci has encountered difficulties in terms of computational costs and interpretability of results. On the computational side, progress has been made using approaches based on convex optimization such as the lasso [14], developing accurate methods to screen variables [15–17], and relying on variational Bayes [18, 19]. There are, however, remaining challenges. Firstly, the genetics community is, correctly, very sensitive to the need of replicability, and finite samples guarantees for the selected variants are sought. Unfortunately, this has been difficult to achieve with techniques such as the lasso: [20] attempts to use stability selection, [21] does a simulation study of a variety of penalized methods, showing that tuning parameters play a crucial role and that standard selection methods for these do not work well, and [22] proposes some analytical approximation of FDR as an alternative to the lasso. Our recent work [23] also explores alternative penalty functions that under some circumstances guarantee FDR control. Secondly, multivariate models encounter difficulties in dealing with correlated predictors, in that the selection among these is often arbitrary: this is challenging in the context of GWAS, when typically there is a substantial dependence between SNPs in the same genetic region.

The suggestion of controlling FDR rather then FWER in genetic mapping studies that expect to uncover a large number of loci was put forward over a decade ago [24–26] and is accepted in the expression quantitative trait loci (eQTL) community, where FDR is the standard error measure. The existence of strong local dependence between SNPs has also posed challenges for FDR controlling procedures. While the Benjamini-Hochberg [10] procedure (BH) might be robust to the correlation between tests that one observes in GWAS, the fact that the same biological association may be reflected in multiple closely located SNPs complicates both the definition and the counting of discoveries, so that it is not immediately evident how FDR should be defined. Prior works [27–29] underscore this problem and suggest solution for specific settings.

This paper proposes a phenotype-aware selective strategy to analyze GWAS data which enables precise FDR control and facilitates the application of multivariate regression methodology, by reducing the dependency between the SNPs included in final testing. The Methods section starts by briefly recapitulating the characteristics of GWAS, with reference to an appropriate count of discoveries and the identification of a meaningful FDR to control. We introduce our selective strategy and provide some general conditions under which it controls the target FDR. We then describe a specific selection procedure for GWAS analysis and describe how it can be coupled with standard BH for univariate tests, or with SLOPE [23] to fit multivariate regression. In the Results section we explore the performance of the proposed methodology with simulations and analyze a dataset collected in the study of the genetic basis of blood lipids. In both cases, the FDR-controlling procedures we propose allow us to explain a larger portion of the phenotype variability, without a substantial cost in terms of increased false discoveries.

## 1 Methods

### 1.1 The GWAS design, dependence and definition of discoveries

The goal of a GWAS study is to identify locations in the genome that harbor variability which influences the phenotype of interest. This is achieved using a sample of *n* individuals, for whom one acquires trait values *y_i_* and genotypes at a collection of *M* SNPs that span the genome.

Following standard practice, we summarize genotypes by the count of copies of minor allele that each individual has at each site, resulting in a *n* × *M* matrix *X*, with entries *X_ji_* ∈ {0,1,2}. The variant index *j* is taken to correspond to the order of the position of each SNP in the genome. The true relation between genetic variants and phenotypes can be quite complex. For simplicity, and in agreement with the literature, we assume a linear additive model, which postulates that the phenotype value *y_i_* of subject *i* depends linearly on her/his allele counts at an unknown set *С* of causal variants. Since there is no guarantee a priori that the variants in *С* are part of the genotype set, we indicate their allele counts with *Z_i_j*, letting 
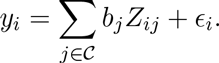

Investigating the relation between *y* and *X* is helpful to learning information about the set of causal variants *С* and their effects *b_j_* in two ways: (1) it is possible that some of the causal variants are actually genotyped, so that *Z_ij_* = *X_ik_* for some *k*; (2) most importantly, the set of *M* genotyped SNP contains reasonable proxies for the variants in *С*. To satisfy (2), GWAS are designed to capitalize on the local dependence between variable sites in the genome known as linkage disequilibrium (LD), which originates from the modality of transmission of chromosomes from parents to children, with modest recombination. The set of *M* genotyped SNPs is chosen with some redundancy, so that the correlation between *X_j_* and *X_j+k_* is expected to be non-zero for *k* in a certain range: this is to ensure that any non-typed casual variant *Z_l_* will be appreciably correlated with one (or more) of the typed *X_j_*s which are located in the same genomic region. Any discovered association between a SNP *X_j_* and the phenotype *y* is interpreted as an association between *y* and *some variant* in the genomic *neighborhood* of *X_j_*. This design has a number of implications for statistical analysis:

1. Often, the existence of an association between *y* and each typed variant *X_j_* is queried via a test statistic *t_j_* which is a function of *y* and *X_j_* only: these test statistics are “locally” dependent, with consequences for the choice of multiple comparison adjustment, that, for example, might not need to be as stringent as in the case of independence.
2. When multivariate regression models are used to investigate the relation between *y* and *X*, one encounters difficulties due to the correlation between regressors—the choice among which is somewhat arbitrary.
3. The fact that the true causal variants are not necessarily included among the genotyped SNPs makes the definition of a true/false association non-trivial.

We want to underscore the last point. To be concrete, let's assume the role of each variant *X_j_* is examined with *t_j_*, the t-statistic for 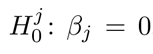, with (*β_j_* defined in the univariate regression *y_i_* = *α* + *β_j_ X_ij_* + *∈_i_*. Even if none of the *M* genotyped variants are causal, a number of them will have a coefficient *β_j_* ≠ 0 in these reduced models: whenever *X_j_* is correlated with one of the variants in *С*, 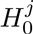 should be rejected. Indeed, simulation studies that investigate the power and global error control of different statistical approaches routinely adopt definitions of “true positive” that account for correlation between the known causal variant and the genotyped SNPs (see [21] for a recent example). At the same time, a rejection of 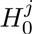 should not be interpreted as evidence of a causal role for *X_j_*: in fact, geneticists equate discovery with the identification of a genomic location rather then with the identification of a variant. The rejection of 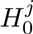 for a number of correlated neighboring SNPs in a GWAS is described in terms of the discovery of one single locus associated with the trait of interest. The number of reported discoveries, then, corresponds to the number of distinct genomic regions (whose variants are uncorrelated) where an association has been established. This discrepancy between the number of rejected hypotheses and the number of discoveries has important implications for FDR controlling strategies, which have received only a modest attention in the literature. Siegmund and Zhang [29] suggest that in situations similar to those of GWAS, neighboring rejections should be grouped and counted as a single rejection and that the global error of interest should be the expected value of the “proportion of clusters that are falsely declared among all declared clusters”. This FDR of clusters—a notion first introduced in [28]—is not the error rate controlled by the Benjamini-Hochberg [10] procedure on the p-values for the 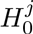 hypotheses. Indeed, because FDR is the expected value of the ratio of the random number of discoveries, its control depends crucially on how one decides to count discoveries. In [30] we give another example of how controlling FDR for a collection of hypotheses does not extend to controlling FDR for a smaller group hypotheses logically derived from the initial set. Both in the setting described here and in [30], targeting FWER would have resulted in less surprising behavior: assuring that the probability of rejecting at least one null 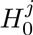 is smaller than a level *α* would also guarantee that the probability of rejecting a null cluster of hypotheses is smaller than *α*. Siegmund and Zhang [29] study a setting that is close to our problem and propose a methodology to control their target FDR relying on a Poisson process distribution for the number of false discoveries. We investigate here a different approach: one that is more tightly linked to the GWAS design, is adapted to the variable extent of LD across the genome, and capitalizes on results in selective inference [31].

### 1.2 Controlling the FDR of interesting discoveries by selecting hypotheses

The approach we study emerged from our interest in using multivariate linear models to analyze the relation between *y* and *X*, so it is useful to motivate it in this context. Suppose both *X_j_* and *X_j+1_* are strongly correlated with the untyped causal variant *Z_k_*. When univariate regression is used as the analysis strategy, both the test statistics *t_j_* and *t_j+1_* would have large values, resulting in the discovery of this locus. Instead, the marginal p-values for the coefficients of *X_j_* and *X_j+1_* derived from a multivariate model that includes both would be large; and model selection strategies would rather arbitrarily lead to the inclusion of one or the other regressor, leading to an underestimate of their importance when resampling methods are used to evaluate significance. If using multivariate linear models, one would achieve the best performance if, from the start, only one of *X_j_* and *X_j+1_* (the most strongly correlated with *Z_k_*) is included among the possible regressors. A natural strategy is to prune the set of *M* typed SNPs to obtain a subset of *m* quasi-orthogonal ones and supply these to the model selection procedure of choice. However, this encounters the difficulty that the best proxy for some of the causal variants might have been pruned, resulting in a loss of power. It seems that ideally one would select from a group of correlated SNPs the one that has the strongest correlation with the trait to include among the potential regressors. Unfortunately, this initial screening for association would invalidate any guarantees of the model selection strategy, which operates now not on *m* variables, but on *m selected* ones. The emerging literature of selective inference, however, suggests that we might be able to appropriately account for this initial selection step, preserving guarantees on error rate control.

Abstracting from the specifics of multivariate regression, consider the setting where a collection ℋ of *M* hyspotheses 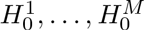 with some redundancy is tested to uncover an underlying structure of interest. The hypotheses in *ℋ* can be organized linearly or spatially and are chosen because a priori they provide a convenient and general way of probing the structure; however, it is expected that a large portion of these will be true, and that when one 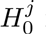 is false, a number of neighboring ones would be also false. In case of GWAS, these clusters of false hypotheses would correspond to markers correlated with causal mutations. Because of the mismatch between *ℋ* and the underlying structure, the number of scientifically interesting discoveries does not correspond to the number of rejected 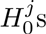 and strategies that control the FDR defined in terms of these might not lead to satisfactory inference. Specifically, as noted in [29], “a possibly large number of correct rejections at some location can inflate the denominator in the definition of false discovery rate, hence artificially creating a small false discovery rate, and lowering the barrier to possible false detections at distant locations”. This problem was recognized already in [27] and [28], who introduce the notion of cluster FDR and suggest defining *apriori* clusters of hypotheses, corresponding to signals of interest and apply FDR controlling strategies to hypotheses relative to these clusters. We take here a different approach, where “clusters” of hypotheses are defined *after looking at the data*, and used to select a subset of repressentative hypotheses. Only this subset is then tested, with a procedure that accounts for this initial selection.

Formally, let *y* indicate the data used to test the hypotheses in *ℋ* and let *S*(*y*) be a selection procedure that, on the basis of the data, identifies a subset *ℋ^s^* of s representative hypotheses. Let 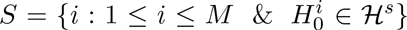 be the set of their indexes, so that it is relevant to control the following FDR_s_: 
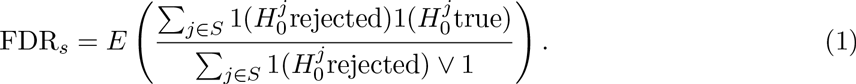

In other words, the decision of acceptance/rejection is made only for the hypotheses in the selected set. The work of [32] and [31] suggests a possible strategy to control FDR_s_ at level q: apply BH to the p-values *p*[_S_] corresponding to the subset of hypotheses *ℋ^s^*, targeting the more stringent level *q*|*S*|/*M* to penalize for the initial selection. According to this strategy, the smallest p-value *p*_[*s*](1)_ for *ℋ^s^* would be compared to |*S*|*q*/*M* × 1/|*S*| = *q*/*M*, and *P*[_*S*](*i*)_ would be compared to *qi*/*M*: the p-value thresholds are identical to those implied by BH on *ℋ*, but the number of hypotheses tested is smaller and the hypotheses are more clearly separated. This prevents the excessive deflation of the BH threshold that results when each true discovery is represented by many rejected hypotheses, and therefore helps to control the number of false discoveries.

The results in [31] imply that if *S*(y) is a simple selection rule, the procedure described above controls the selective FDR 
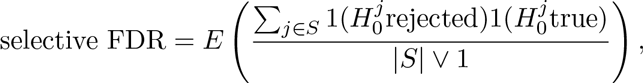
whenever BH applied to *ℋ* would control the standard FDR, 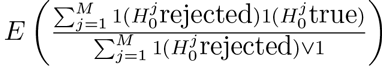. If the selection is stringent enough, controlling the selective FDR might be meaningful. Building on the results obtained in [33], we can also prove that for a general class of selection rules the same procedure leads to control of FDR*s* at level *q*, as long as the distribution of p-values follows the condition of positive regression dependence on a subset (PRDS), described in [34]. As noted in [25], PRDS condition can be loosely interpreted as the requirement of the positive correlation between p-values at linked markers, and in this way it corresponds well to the practice of GWAS.

#### Theorem 1

*FDR control for selected hypotheses*. *Let S*(*y*) *be a selection procedure*, *and let R^S^ be the number of rejections derived by applying BH with target q*|*S*|/*M on the selected hypotheses ℋ^S^*. *If the p-values are PRDS and the selection procedure is such that R^S^*(*p*_1_,… ,*p_M_*) *is non-increasing in each of the p-values p_i_*, *rejecting R^S^ guarantees control of* FDR_*s*_.

*Proof*. Letting *ℋ*_0_ be the collection of true null hypotheses in *ℋ* and 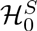 the set of true null hypotheses in *ℋ^S^*, we write FDR_*s*_ as 
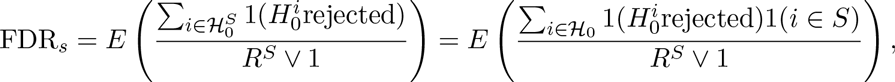
where *R^S^* indicates the number of rejections resulting from applying the BH rule with target level *q*|*S*|/*M* to the p-values *p*[_*S*_] of *ℋ^S^*. Going forward, we write *R^S^* instead of (*R^S^* ⋁ 1) for simplicity. Recalling that a hypothesis is rejected if its p-value is smaller than the BH threshold, exchanging the order of summation and expectation, and multiplying and dividing by *q*/*M* we have 
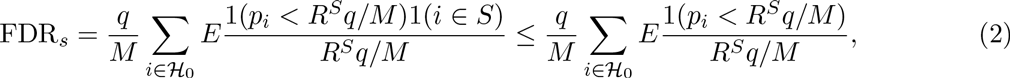
where the last inequality comes from relaxing a restriction. We now recall Lemma 1 from [33], which states that for a set of p-values that satisfy PRDS, when *f*:(*p*_1_,… ,*p_M_*) ⤒[0,1] is non-increasing, 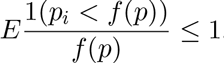. Under the assumption that *f*(*p*_1_,… ,*p_M_*):= *R^S^ q*/*M* is non-increasing we then have our result, as 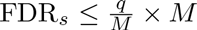.

An example selection procedure that satisfies the assumptions of the theorem is as follows: the hypotheses *ℋ* are separated in groups a priori and from each group, *S(y)* selects the hypothesis with the smallest associated p-value. In the next section, we describe a slightly more complicated selection procedure *S*(*y*), that appears appropriate for the case of GWAS, and where the separation of hypotheses into groups is data-driven. While this procedure may not satisfy the assumption that the number of rejections is a non-increasing function of the p-values, our extensive simulations studies suggests that its use in the context of Theorem 1 still leads to FDR, control.

### 1.3 A GWAS selection procedure: phenotype-aware cluster representatives

In the context of genetic association studies, the selection function *S*(*y*) defined in procedure 1 and illustrated in Figure 1 emerges quite naturally. One starts by evaluating the marginal association of each SNP to the phenotype using the p-value of the t-test for its coefficient in a univariate regression. Then, SNPs with a p-value larger than threshold *π* are removed from consideration. The collection of remaining SNPs is further pruned to obtain a selected set *S* with low correlation, so that each variant *X_i_* ∈ *S* can be equated to a separate discovery. To achieve this, we define clusters of SNPs using their empirical correlation in our sample, starting from the variants with the strongest association to the phenotype, which are selected as cluster representatives.

**Figure 1.**
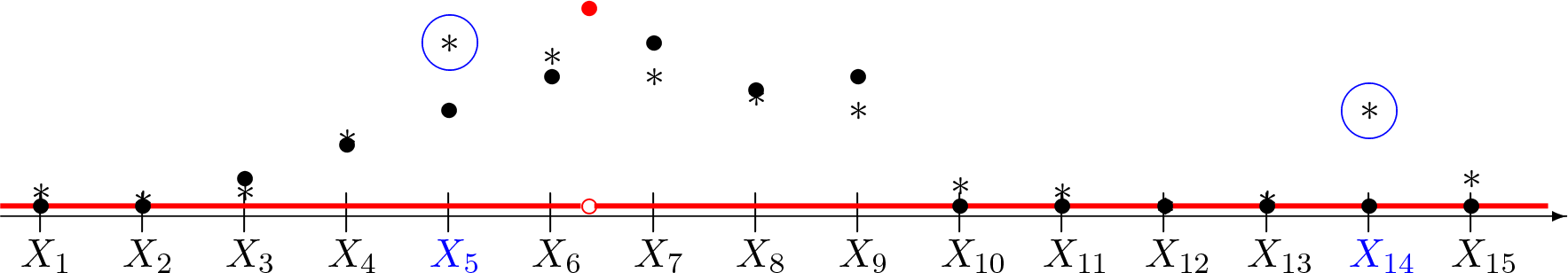
Phenotype-aware cluster representatives. *The x-axis represents the genome, with the locations of genotyped SNPs X_i_ indicated by tick-marks. The true causal effect of each position of the genome is indicated in red: there is only one causal variant in this region, between SNPs X_6_ and X_7_. Solid black circles indicate the value of β_i_, coefficient of X_i_ in a linear approximation of the conditional expectation E*(*y*|*X_i_*). *Asterisks mark the estimated (μ_i_s in the sample. The SNPs X_5_ and X_14_, selected as cluster representatives in this schematic diagramm, are indicated in blue*.

**Procedure 1.**
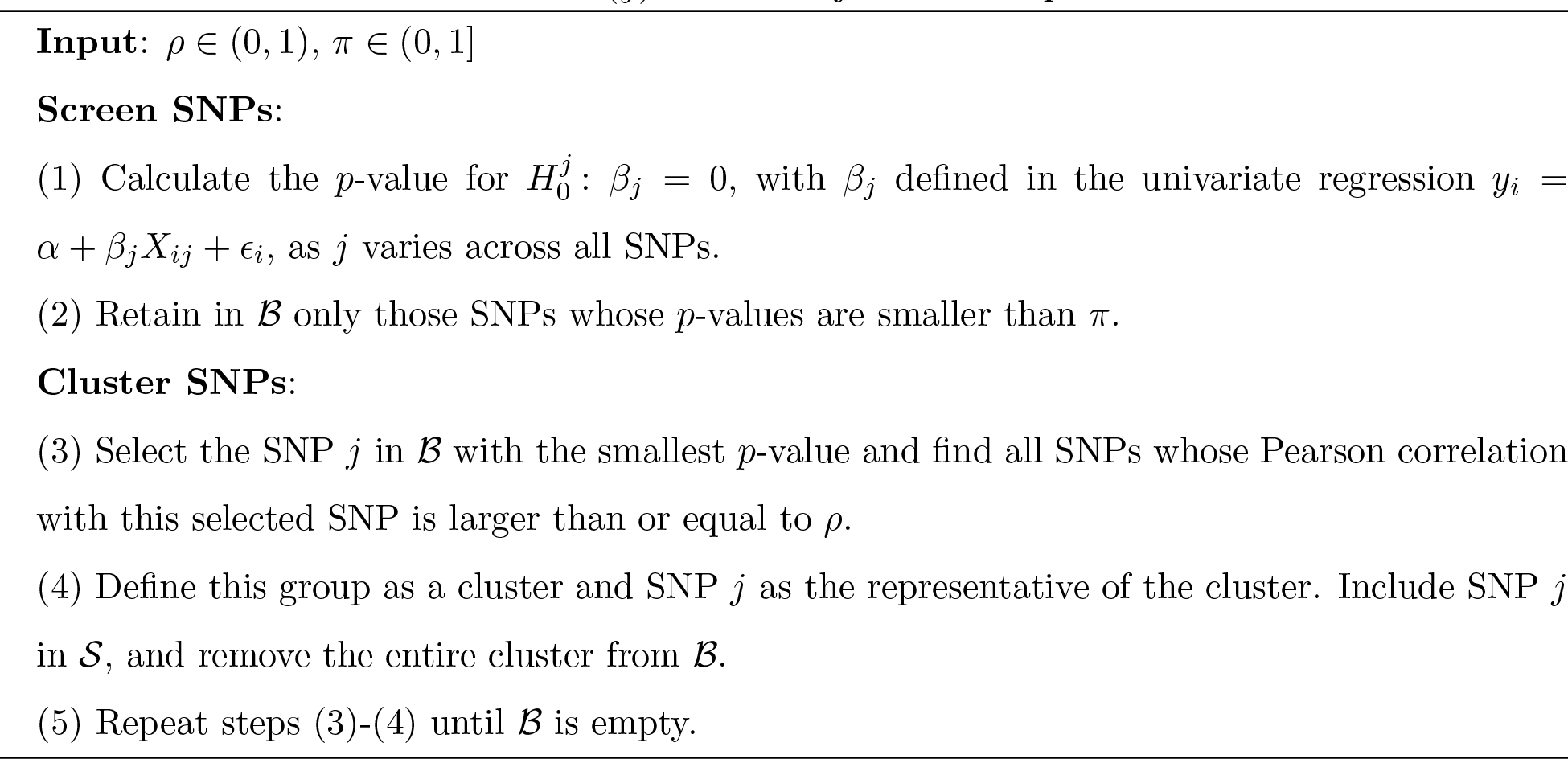
Selection function *S*(*y*) to identify cluster representatives

Procedure 1 has two tuning parameters: *π* and *ρ*, corresponding to the two steps of the selection. The screening in steps (1)-(2) is similar to that described in [13, 15] for model selection procedures, where the parameter *π* controls the stringency of the selection based on univariate association. Its influence is minimal if marginal tests are used to control FWER. However, large values of *π* result in a larger dimensions of selected clusters, leading to increased number of false discoveries when controlling FDR. On the other hand, in the context of multivariate regression, it is possible to uncover a role for variants that have weak marginal effects due to masking: to enable this, one must not be too stringent in the initial screening step. In all the simulations and data analyses presented here we have used *π* = 0.05, which seems to be a good compromise. The results in [13, 15] can provide additional guidance on the choice of *π*.

Steps (3)-(5) of procedure 1 aim to “thin” the set of SNPs on account of the dependency among them. This is related to the selection of tag SNPs [35], for which there is an extensive literature, and is similar to correlation reduction approaches [36]. A defining characteristic of procedure 1, however, is that both the SNP clusters and their representatives are selected with reference to the phenotype of interest. This ensures that the representatives maximize power, and that the location of the true signal is as close as possible to the center of the respective cluster. This also reduces the probability of the selection of more than one SNP per causal variant. The value of *ρ* needs to be set with reference to the sample size and the density of the available markers. Indeed, we suggest that researchers run a simple simulation (as in the one described in the first part of the Results section) to select appropriate values for *ρ* (see Discussion section for further remarks on this). Certainly, procedure 1 is but one possibility for creating clusters. For example, one might want to include information on physical distance in the formation of clusters. In our experiments, however, this has not led to better performance.

We now consider two approaches to the analysis of GWAS data that can be adopted in conjunction with the selection of cluster representatives to control the FDR_*s*_.

### 1.4 Univariate testing procedures after selection

By and large, the most common approach to the analysis of GWAS data relies on univariate tests of association between trait and variants. This has advantages in terms of computational costs, handling of missing data, and portability of results across studies. We therefore start by considering how to control relevant FDR in this context.

While most disease-related GWAS aim to control FWER, FDR has been the global error of choice in eQTL studies, and that literature testifies to some of the challenges encountered, in particular, to difficulties related to dependence across tests and hypotheses (see [30] for a detailed description). Starting from [25], it was observed that BH seems to be able to deal with the type of dependence across test statistics induced by LD. However, the relatedness between hypotheses and the lack of one-to-one correspondence between hypotheses and meaningful scientific discoveries remains a problem. For example, when investigating the genetic basis of variation in gene expression, the authors in [37] change the unit of inference from SNPs to genes, so as to bypass the redundancy due to many SNPs in the same neighborhood. Here we address the problem by inviting the researchers to identify the resolution of discoveries prior to testing, but after having observed the data. We consider two different approaches to obtain the p-values for each of the 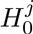 hypotheses: univariate linear regression (which we indicate with SMT for single marker test) and EMMAX [7], a mixed model which allows us to consider polygenic effects. To enable computational scaling, EMMAX only estimates the parameters of the variance component model once rather than for every marker. We use SMTs and EMMAXs to denote the procedures that consist in testing the set of hypotheses *H^s^* corresponding to cluster representatives, using p-values obtained with SMT and EMMAX, respectively, and identifying rejections with the BH_s_ procedure described below.

**Procedure 2.**
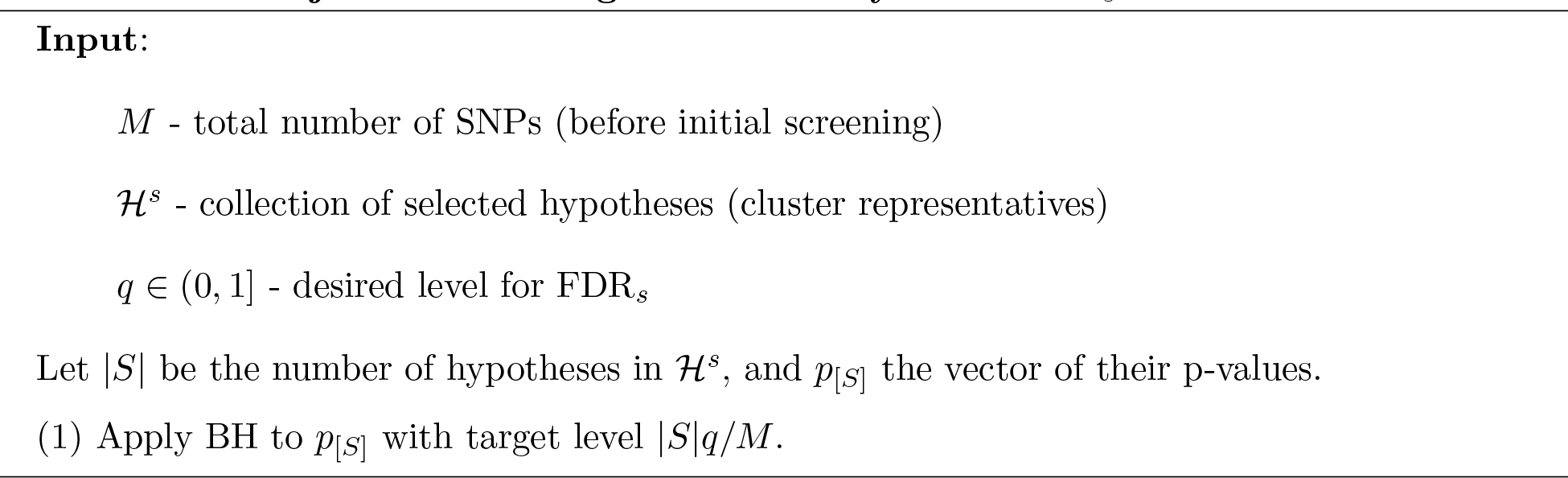
Benjamini-Hochberg on selected hyotheses BH_s_

### 1.5 GeneSLOPE - FDR control in multivariate regression.

SLOPE [23] is a recently introduced extension of the lasso that achieves FDR control on the selection of relevant variables when the design is nearly orthogonal. Specifically, assume the following model 
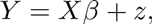
where *X* is the design matrix of the dimension *n* × *M*, *z* ~ *N*(0, σ^2^I_*n*×*n*_) is the *n*-dimensional vector of random errors, and *β* is the M-dimensional vector of regression coefficients, a significant portion of which is assumed to be zero. For a sequence of non-negative and non-increasing numbers *λ*_1_,…, λ_*M*_, the SLOPE estimate of *β* is the solution to a convex optimization problem 
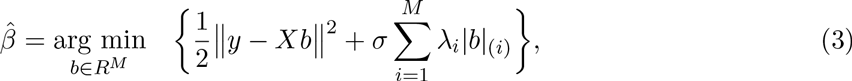
where |*b*|_(1)_ ≤ … ≤ |*b*|_(*M*)_ are sorted absolute values of the coordinates of *b*.

If we define a discovery as *i* such that the estimated 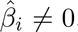, and a false discovery as the case where 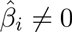 but the true *β_i_* = 0, [23] provides the sequence of *λ_i_* (corresponding to the sequence of decreasing thresholds in BH), which provably controls FDR at a desired level if the design matrix *X* is orthogonal. Moreover, the modified sequence λ—described in procedure 4 in the Appendix— has been shown in simulation studies to achieve FDR control in genetic studies when SNPs are nearly independent and the number of non zero *β*'s is small or moderately large. Note that, as for other shrinkage methods [38, 39], the results of SLOPE depend on the scaling of explanatory variables: the values of the regularizing sequence in procedure 4 assume that explanatory variables are “standardized” to have zero mean and a unit *l_2_* norm. Moreover, since in most cases the variance of the error term σ^2^ is unknown and needs to be estimated, in [23] an iterative procedure for the joint estimation of *σ* and the vector of regression coefficients was proposed. This is described in the Appendix as procedure 5 and follows closely the idea of *scaled lasso* [40]. All these data preprocessing and analysis steps are implemented in R package *SLOPE*, available on CRAN.

The fact that SLOPE comes with finite sample guarantees for the selected parameters makes it an attractive procedure for GWAS analysis. However, the presence of substantial dependence between SNPs (regressors *X_j_*) presents challenges: on the one hand, the FDR-controlling properties have been confirmed so far only when the explanatory variables are quasi-independent; and on the other hand, the definition of FDR is problematic in a setting where the true causal variants are not measured and *X* contains a number of correlated proxies, similarly as for univariate procedures. The identification of a subset of variants with procedure 1 takes care of both aspects: the regressors are not strongly correlated and, for sufficiently small *ρ*, they represent different locations in the genome, so that we can expect the projection of the true model in the space they span to be sparse and the number of 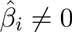 to capture the number of scientifically relevant discoveries. We therefore propose as a potential analysis pipeline the application of procedure 1 followed by procedure 3, which outlines the application of SLOPE to the selected cluster representatives. Both procedures have been implemented in the R package GeneSLOPE, which is available on CRAN and can handle typical GWAS data provided in PLINK format.

**procedure 3.**
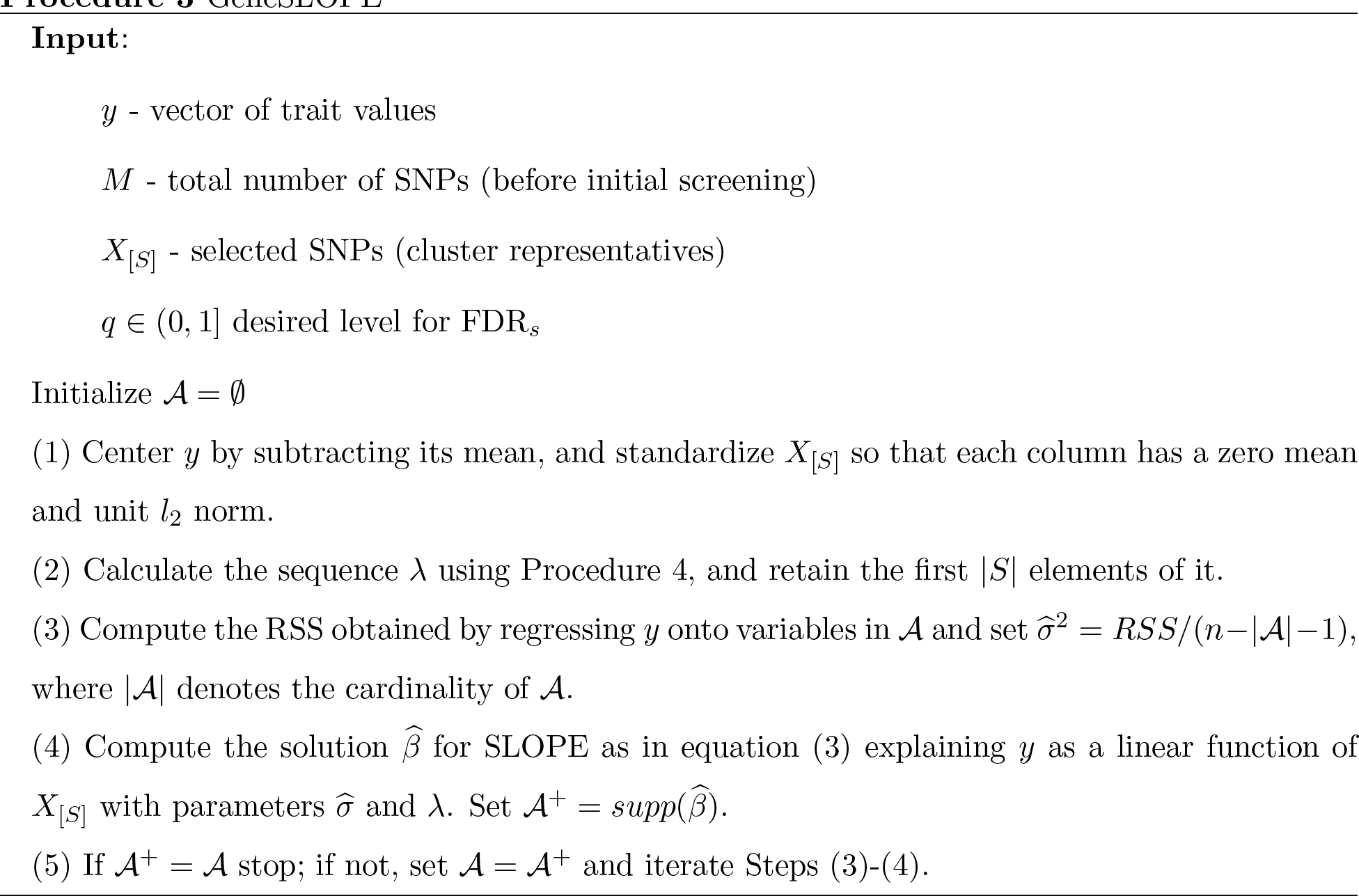
GeneSLOPE

## 2 Results

To test the performance of the proposed algorithms we relied on simulations and real data analysis. In both cases, genotype data came from the North Finland Birth Cohort (NFBC66) study [41] , available in dbGaP under accession number phs000276.v2.p1 (http://www.ncbi.nlm.nih.gov/projects/gap/cgi-bin/study.cgi?study_id=phs000276.v2.pl). The raw genotype matrix contains 364,590 markers for 5,402 subjects. We filtered the data in PLINK to exclude copy number variants and SNPs with Hardy-Weinberg equilibrium *p*-value < 0.0001, minor allele frequency < 0.01, or call rate < 95%. This resulted in an *n* × *M* predictor matrix with *n* = 5,402 and *M* = 334, 103. When applying GeneSLOPE, missing genotype data were imputed as the SNP mean.

For simulations, the trait values are generated using the multiple regression model: 
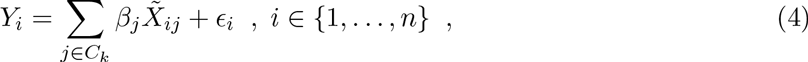
where 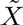 is the standardized matrix of genotypes, *C_k_* is the set of indices corresponding to “causal” mutations and *∈_i_* ~ *N*(0,1). The number of causal mutations takes the value *k* ∈ {20, 50,80,100}, and in each replicate, the *k* “causal” features are selected at random from a subset of the *M* SNPs. For each *k*, the values of *β_j_* are evenly spaced in the interval [SignalMin, SignalMax], with 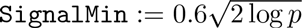 and 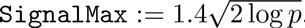. As a result, the smallest genetic effect is rather weak (heritability in a single QTL model *h*^2^ = 0.0017), while the strongest effect is relatively large (*h*^2^ = 0.0091). Each scenario is explored with 100 simulations.

In evaluating FDRs and power, we adopt the following conventions, which we believe mimic closely the expectations of researchers in this field: the null hypothesis relative to a SNP/cluster representative is true if the SNP/cluster representative has a correlation less then 0.3 with any causal variant. Similarly, a causal variant is discovered if at least one of the variants in the rejection set has correlation of at least magnitude 0.3 with it.

In addition to evaluating performance in the context of simulated traits, we apply the proposed procedures to four lipid phenotypes available in NFBC66 [41]: high-density lipoproteins (HDL), low-density lipoproteins (LDL), triglycerides (TG), and total cholesterol (CHOL). We compare the discoveries obtained by the univariate and multivariate procedures on the NFBC data to those reported in [42], a much more powerful study based on 188,577 subjects.

## 2.1 Simulation study

### Cluster sizes

We begin by exploring the distribution of the size of clusters created according to procedure 1. Figure 2 illustrates the size of clusters when the trait was generated according to the model in equation (4) with *k* = 80 and genotypes from the NFBC dataset. We illustrate the results of both the original version of procedure 1 using simple univariate regression (SMT) to obtain the p-values as well as a modification in which the initial p-value calculation is performed using EMMAX. It can be seen that most of the clusters are rather small and do not include more than 5 SNPs. There are no significant differences in the size of clusters created starting from EMMAX or SMT p-values. Of course, differences in the genotype density would result in a differences in the cluster sizes obtained.

**Figure 2:**
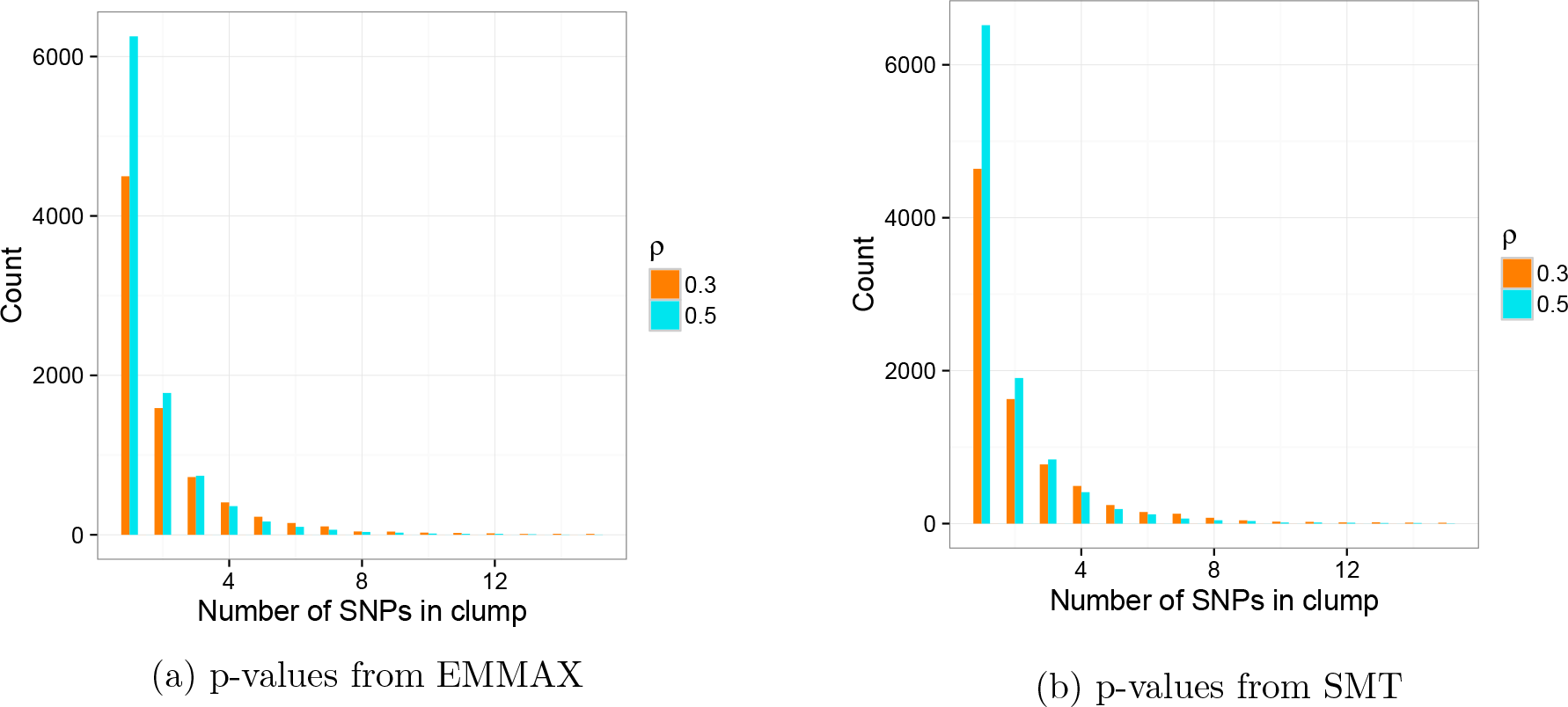
Histograms of the number of SNPs included in each cluster when procedure 1 is applied to p-values calculated from SMT and EMMAX with *π* = 0.05 and *ρ* = 0.3 or *ρ* = 0.5.

### Error control with EMMAX and single marker tests

Figure 3 illustrates the results of simulations exploring the FDR, control properties of BH applied the complete set of *M* p-values obtained from EMMAX or SMT (i.e. with no pre-screening or clustering of the hypotheses) and the corresponding two-step approaches we recommend (EMMAXs and SMTs), where cluster representatives are first chosen using procedure 1 and then discoveries are identified with procedure 2. The FDR, for the traditional version of EMMAX and SMT is calculated mimicking what researchers typically do in practice to interpret GWAS results. Specifically, the SNPs for which the null hypotheses are rejected using BH are supplied to procedure 1 to identify clusters. The realized FDR_*s*_ is defined as the average across 100 iterations of the fraction of falsely selected clusters over all clusters obtained.

Figure 3 illustrates that, in agreement with Theorem 1, EMMAXs controls FDR_*s*_ at all levels of *ρ* and for any number of causal SNPs. In contrast, BH applied to the full set of *p*-values obtained from EMMAX with post-hoc clustering of the discoveries results in a somewhat elevated FDR_*s*_ due to the deflation of the BH threshold. Moreover, EMMAXs offers better control of FDR_*s*_ than SMTs, particularly as the number of causal SNPs increases. This makes sense given that the model assumed by EMMAX is better able to account for polygenic effects than the single-marker test.

**Figure 3:**
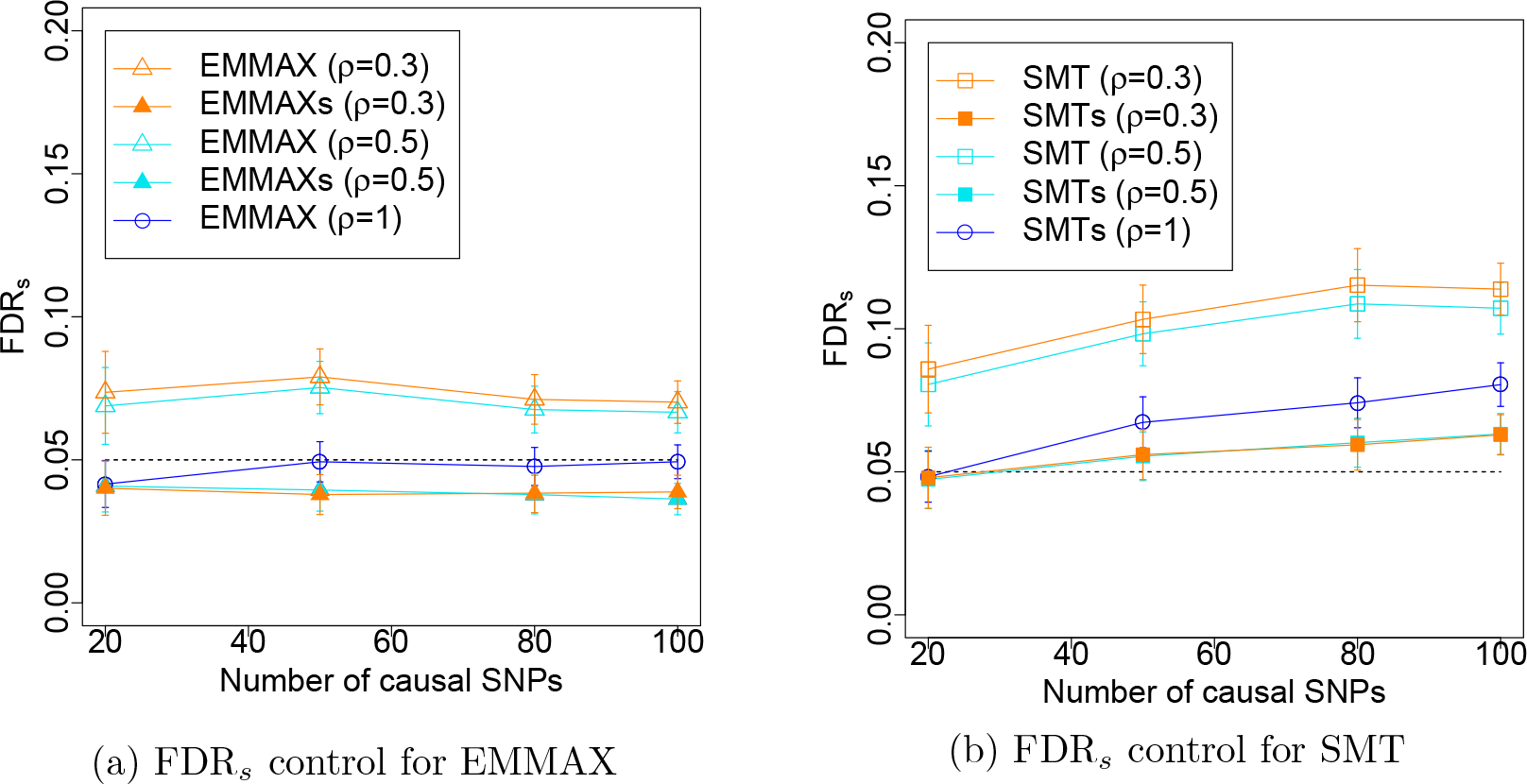
Phenotype-aware cluster representatives. FDR_*s*_ for EMMAX and SMT and the corresponding procedures EMMAXs and SMTs which operate on cluster representatives. The dashed black line represents the target FDR_*s*_ level of 0. 05. Note that EMMAX with *ρ* = 1 (i.e. with no clustering) coincides with EMMAXs, and that the FDR_S_ for this specific case corresponds to the regular FDR. Shapes indicate the procedures: hollow triangles for the application of BH to the collection of *p*-values from EMMAX for all hypotheses followed by clustering of the discoveries, filled triangles for the selective procedure EMMAXs, hollow squares for the application of BH to the collection of *p*-values from EMMAX for all hypotheses followed by clustering of the discoveries, filled squares for the selective procedure SMTs, and hollow circles for the application of BH to the full collection of *p*-values with no clustering. Colors indicate the parameters for clustering: orange for *ρ* = 0.3, turquoise for *ρ* = 0.5, and blue for *ρ* = 1.

### GeneSLOPE error control and power

Figure 4 illustrates the performance of geneSLOPE in terms of FDR_S_ and power in the context of the performance of EMMAXs and SMTs for the same setting and range of k. For all procedures, power decreases as *k* increases, with a slower decay for geneSLOPE. Note that the average power of geneSLOPE is systematically larger than the power of SMTs, with the difference increasing with *k*, while the FDR_*s*_ of geneSLOPE is always smaller then that of SMTs. Figure 4 also demonstrates how using the standard genome-wide significance threshold setting *π* = 5 × 10^−8^ results in a very substantial loss of power as compared to procedures controlling FDR.

**Figure 4:**
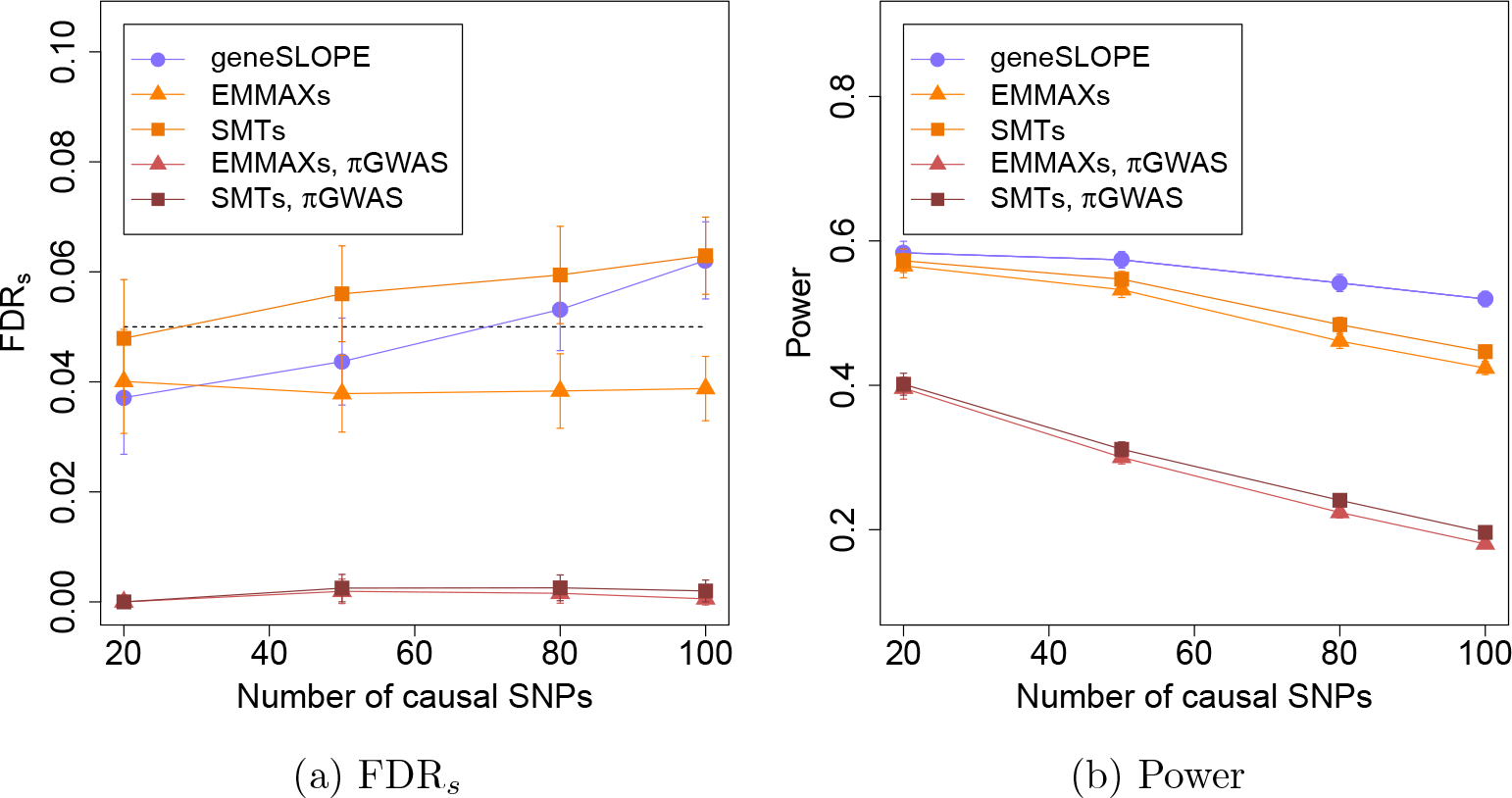
Phenotype-aware cluster representatives. FDR_s_ and power for geneSLOPE when clustering is done with *π* = 0.05, *ρ* = 0.3, and target FDR_*s*_ level 0.05 (marked in slate blue). For comparison, we reproduce from Figure 3 the curves indicating the performance of EMMAXs and SMTs for the same setting (marked in shades of orange). We also include the values of FDR_*s*_ and power when EMMAXs and SMTs are carried out using cluster representatives selected with *π* = 5 × 10^−8^, the standard GWAS genome-wide significance threshold (marked in shades of red). Shapes indicate the procedures: filled circles for geneSLOPE, filled triangles for EMMAXs, and filled squares for SMTs.

#### 2.2 Real Data Analysis

To analyze the lipid phenotypes, we adopted the same protocol described in [41]: subjects that had not fasted or were being treated for diabetes (*n* = 487) were excluded, leaving a set of 4,915 subjects for further analysis. All phenotypes were adjusted for sex, pregnancy, oral contraceptive use, and population structure as captured by the first 5 genotype principal components (computed using Eigensoft [43]); the residuals were used as the trait values *Y_i_* in the subsequent association analysis.

We compare the results of geneSLOPE, EMMAXs and classically applied EMMAX. GeneSLOPE (procedure 1 followed by procedure 3) was applied using *π* = 0.05, *ρ* = 0.3 or 0.5, and *q* = 0.05 or 0.1 (for a total of 4 versions) to a centered and normalized version of the genotype matrix where each column has mean 0 and *l*_2_ norm 1. EMMAXs (procedure 1 followed by procedure 2) was applied with *π* = 0.05, *ρ* = 0.3 or 0.5, and *q* = 0.05 or 0.1. To mimic the standard GWAS analysis, we ran EMMAX identifying as significant those SNPs with p-value ≤ 5 × 10^−8^; to obtain comparable numbers of discovered SNPs we applied procedure 1 to cluster the results.

We compare the discoveries of these three methods on the NFBC data to those reported in [41] , a much more powerful study based on 188,577 subjects. We compute the realized selected false discovery proportion FDP, for each method assuming that SNPs within 1Mb of a discovery (defined as *p* < 5 × 10^−8^) in the comparison study are true positives (even if, of course, the biological truth for the given study population is not known, and the association statistics in [42] are based on univariate tests and may therefore not fully capture the genetic underpinnings of these complex traits). We also seek to understand what proportion of the trait heritability is captured by the selected SNPs: to this end, we estimate the proportion of phenotypic variance explained by the set of genome-wide autosomal SNPs using GCTA [44], and compare this to the adjusted r^2^ obtained from a multiple regression model including the selected cluster representatives as predictors.

The estimated proportion of phenotypic variance explained by genome-wide SNPs is 0.34, 0.32, 0.10, and 0.29 for HDL, LDL, TG and CHOL, respectively. A comparison of the number of discoveries (i.e. the number of selected cluster representatives), number of true discoveries, FDP_*s*_, and r^2^ across methods is given in Figure 5. As an illustrative example, geneSLOPE selections with *π* = 0.05, *q* = 0.1 and *ρ* = 0.5 are shown in Figure 6 along with *p*-values obtained using EMMAX and those obtained in the more highly-powered comparison study [42].

The application on real data illustrates how FDR_*s*_, controlling procedures are more powerful than the standard practice of identifying significant SNPs using a p-value threshold of 5 × 10^−8^. Both EMMAXs and geneSLOPE attain realized selected false discovery proportions that are consistent with the nominal targeted FDR,. There does not appear to be an advantage of multivariate analysis (geneSLOPE) over univariate tests (EMMAXs) in this example: this is consistent with the results in our simulations, which indicate that multivariate analysis is really more powerful when there are many (detectable) signals contributing to the phenotype. While it is by now established that hundreds of different loci contribute to lipid levels, the signal strength in our dataset (which has a modest sample size) is such that only a handful can be identified: in this regime we find no evidence of an advantage for the multivariate linear model.

### 4 Discussion

Following up on an initial suggestion of [29] and reflecting elements of the standard practice, we argue that discoveries in a GWAS study should not be counted in terms of the number of SNPs for which the hypothesis of no association is rejected, but in terms of the number of “clusters” of such SNPs. We propose a strategy to control the FDR of these discoveries that consists in identifying groups of hypotheses on the basis of the observed data, selecting a representative for each group, and applying a modified FDR-controlling procedure to the p-values for the selected hypotheses. We present two articulations of this strategy: in one case we rely on marginal tests of association and modify the target rate of BH on the selected hypotheses; in the other case we build on our previous work on SLOPE to fit a multivariate regression model. We show with simulations and real data analysis that the suggested approaches appear to control FDR_*s*_ and allow an increase in power with respect to the standard analysis methods for GWAS.

**Figure 5:**
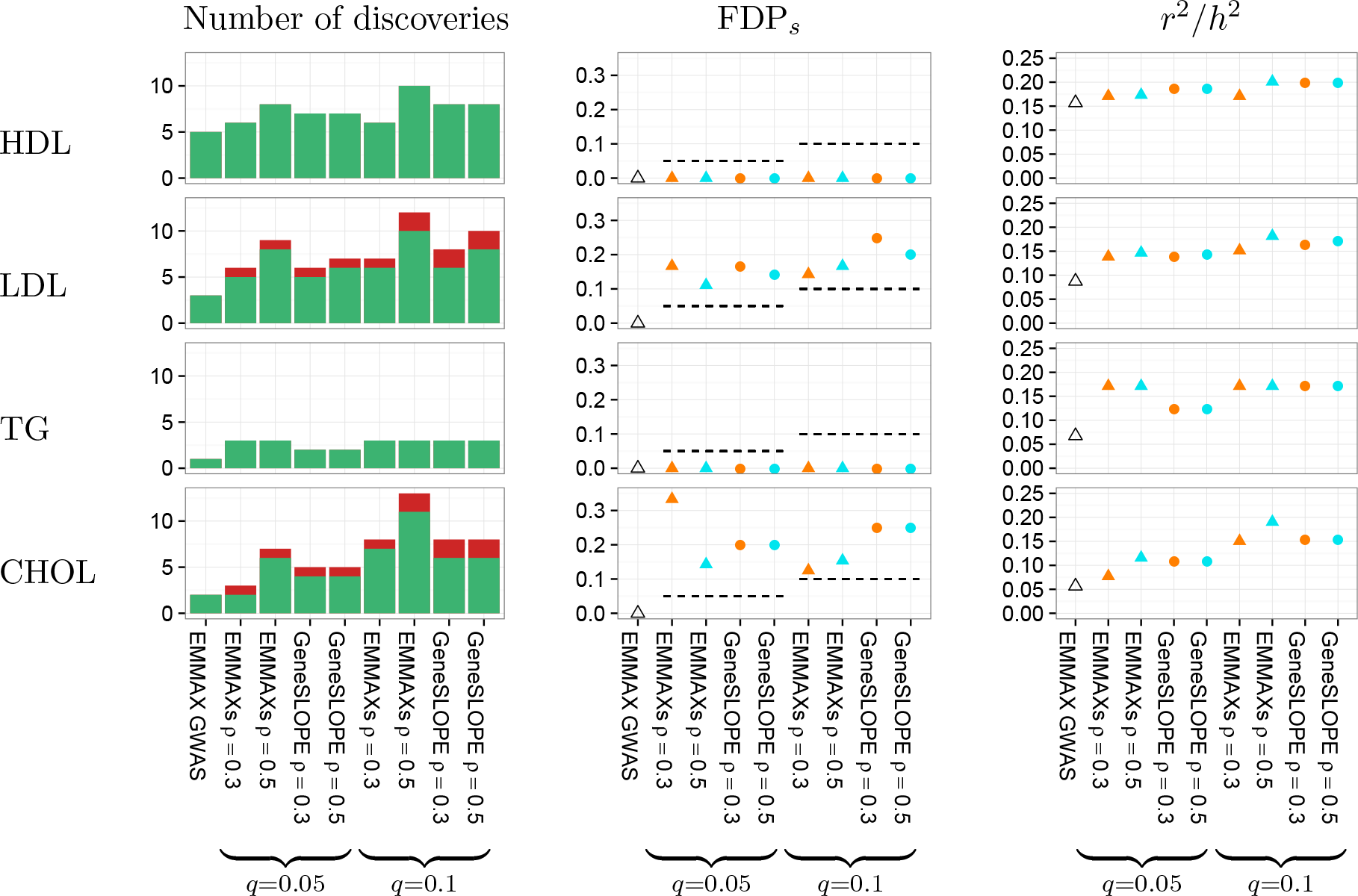
Phenotype-aware cluster representatives. Comparison of methods on the phenotypes high-density lipoproteins (HDL), low-density lipoproteins (LDL), triglycerides (TG), and total cholesterol (CHOL). “Total discoveries” corresponds to the number of selected cluster representatives under each method; in the plot, true discoveries (those within 1Mb of a discovery in [42]) are marked in green, while false discoveries (those not within 1Mb of a discovery in [42]) are marked in red. FDP_S_ is the realized selected false discovery proportion, and *r*^2^/*h*^2^ is the adjusted *r*^2^ obtained when using the set of selected cluster representatives as predictors in a multiple regression model divided by the proportion of phenotype variance explained by genome-wide SNPs obtained using GCTA.

The idea of identifying groups of hypotheses and somehow transferring the burden of FDR control from the single hypothesis level to the group one is not new [27, 28]. In particular, two recent contributions to the literature can be considered parallel to our suggestions. In the context of tests for marginal association, Foyel-Barber and Ramdas [33] propose a methodology to control FDR both at the level of single hypotheses and groups. In the context of multivariate regression, [45] extends SLOPE to control the FDR for the discoveries of groups of predictors. Both these contributions, however, are substantially different from ours in that they require a definition of groups prior to observation of the data. Instead, our “clusters” are adaptive to the signal, and identified starting from the data. This assures that the group of hypotheses are centered around the locations with strongest signal.

Defining cluster representatives that are input to a multivariate regression framework allows us to think more carefully about what FDR means in the context of a regression model that does not include among the regressor the true causal variants, where one is substantially looking for relevant proxies. In their recent work [46], Foyel-Barber and Candes take a different approach, deciding to focus on the directional FDR. The knock-off filter provides an attractive methodology to analyze GWAS data. However, it still requires an initial selection step: top performance can be achieved only when the selected features are optimally capturing the signal present in a given dataset. We believe that the cluster representatives approach has a substantial edge at this level over, for example, running LASSO with only a modest penalization parameter.

We consider here a fairly simple strategy to construct clusters of SNPs, exploring two possible levels of resolution, corresponding to *ρ* = 0.3 and *ρ* = 0.5. In reality, depending on sample size and genotype density, each dataset might have a different achievable level of resolution. The study of how this can be adaptively learned is deferred to future work.

It should be noted while we conduct formal testing only on the selected set of cluster representatives, when the null hypothesis of no association is rejected for a selected SNP, the entire cluster is implicated. In other words, in follow-up studies, the entire region spanned by the cluster should be considered associated with the trait in question.

Finally, we would like to underscore how, even if we have here focused on the case of GWAS, adopting a selective approach might have wide range applications whenever there is not an exact correspondence between the hypotheses conveniently tested and the granularity of the scientific discoveries. Further studies of the emerging literature on selective inference should lead to better understanding of the theoretical properties of the method we propose as well as to the identification of other possible strategies.

**procedure 4.**
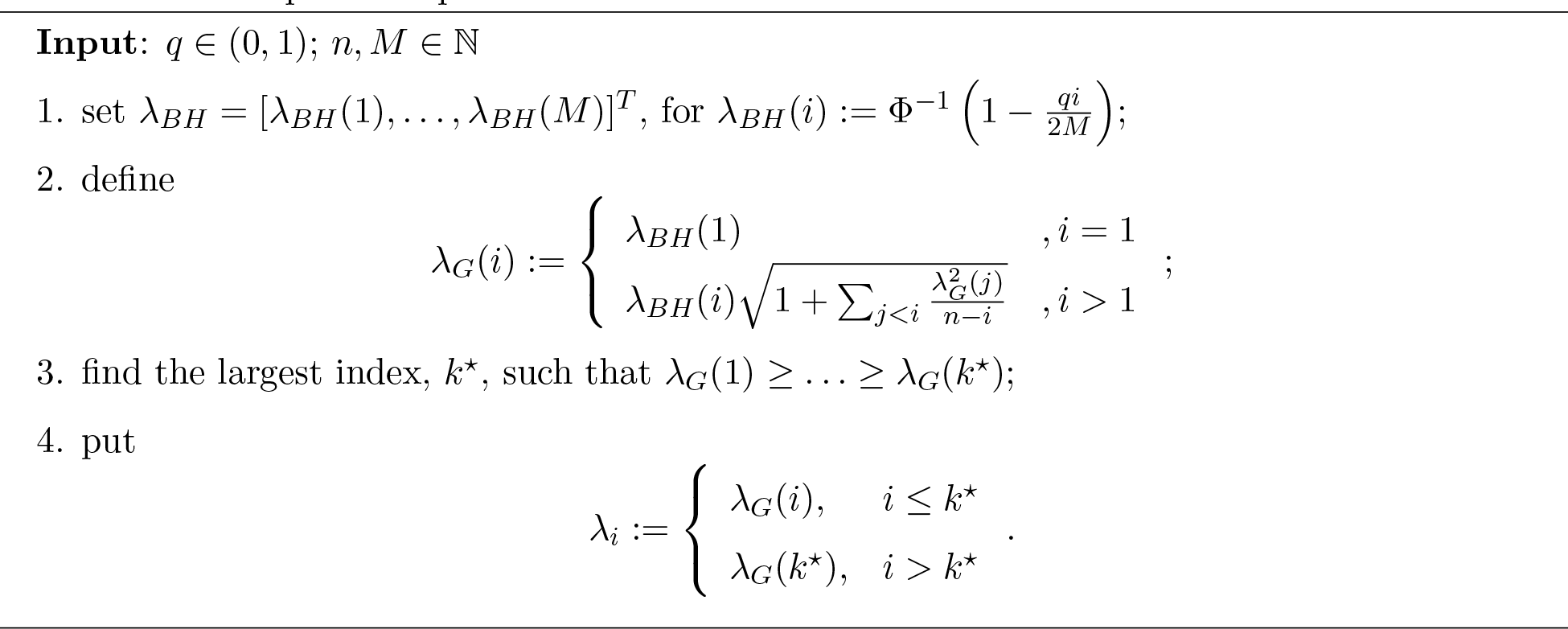
Sequence of penalties λ for SLOPE

**procedure 5.**
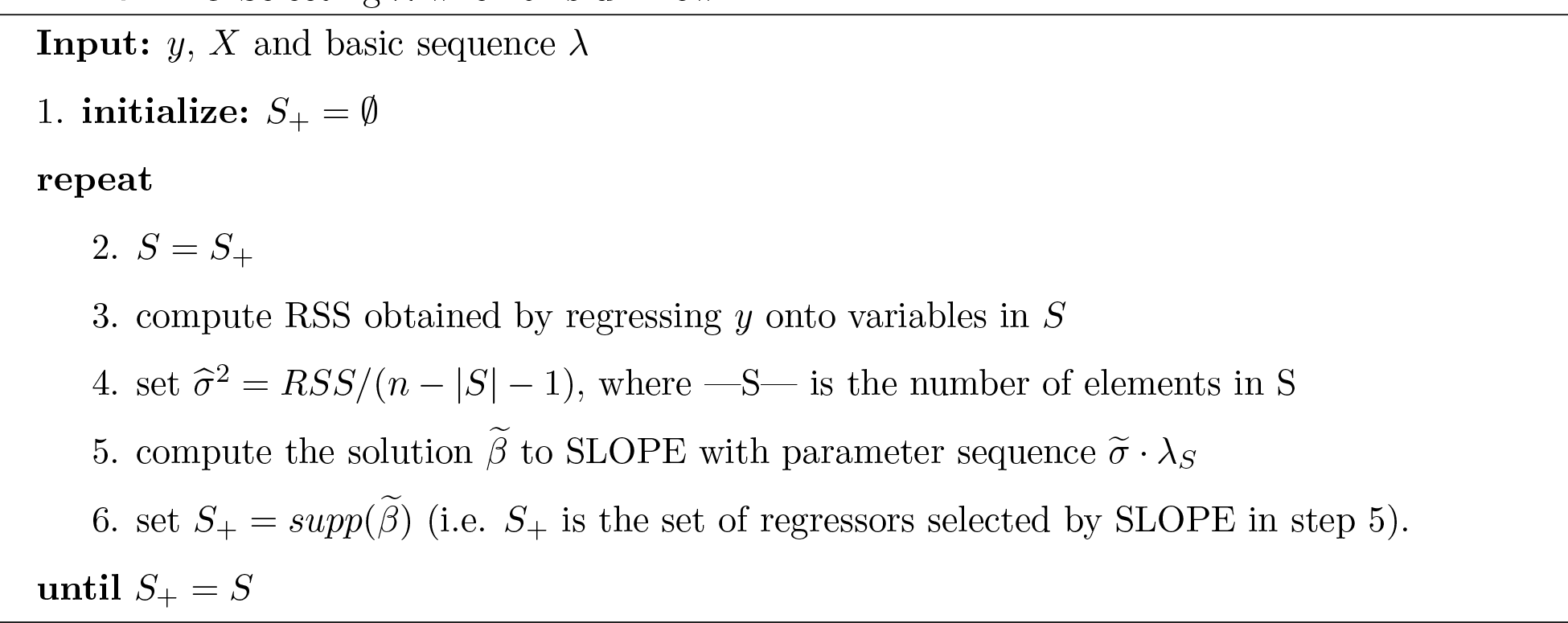
Selecting λ when σ is unknown

**Figure 6:**
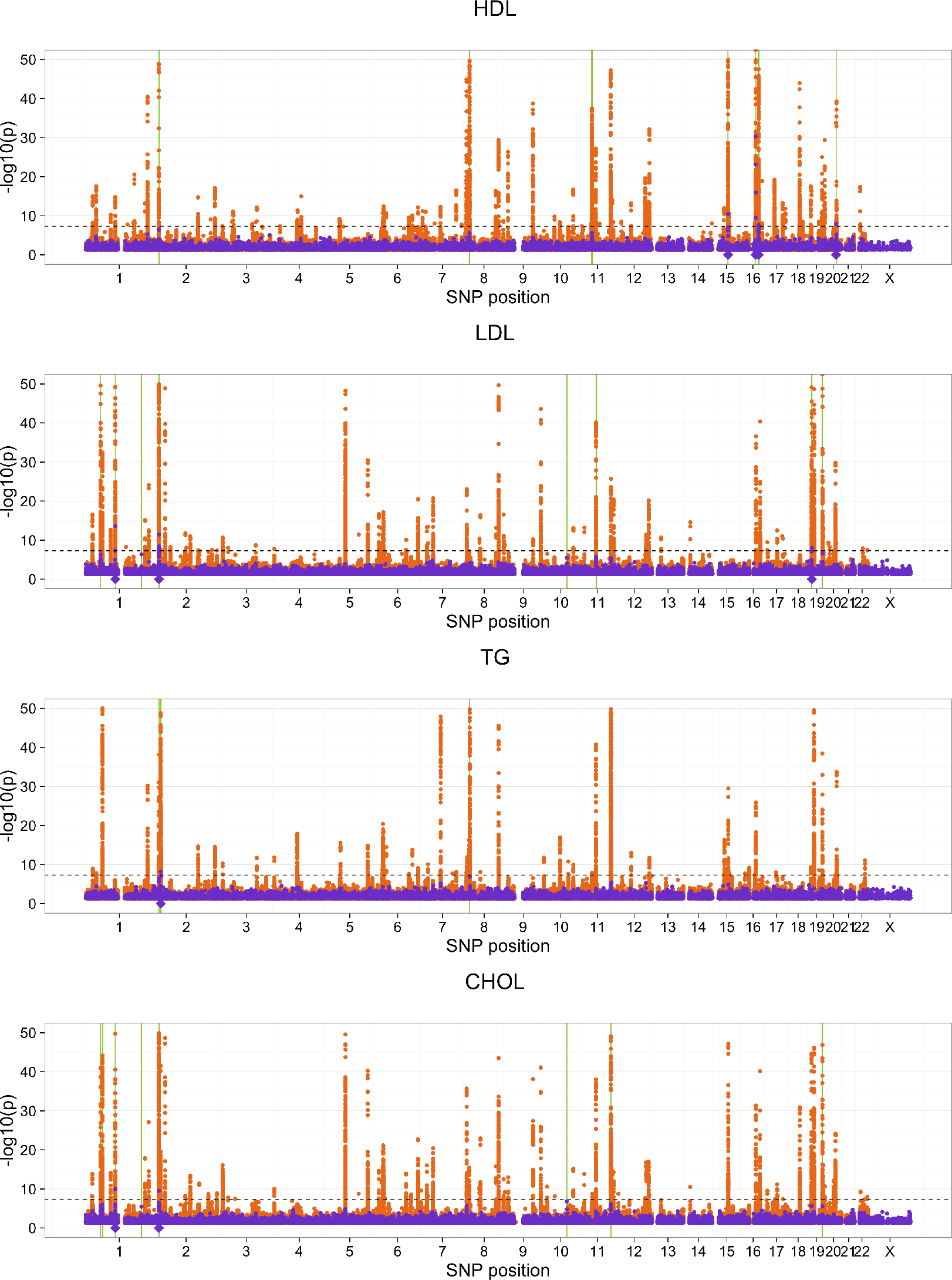
Phenotype-aware cluster representatives. GeneSLOPE selections using *π* = 0:05, *ρ* = 0.5, and target FDR_*s*_ 0.1 are marked using solid green bars for cluster representatives and semi-transparent bars for the remaining members of the cluster. *P*-values from EMMAX (purple) and the Global Lipids Genetics Consortium comparison study (orange) are plotted on the −log10 scale. The horizontal dashed line marks a significance cut-off of 5 × 10^−8^, and the purple diamonds below the x-axis represent selected cluster representatives under EMMAX using *π* = 0:05, *ρ* = 0:3, and a *p*-value threshold of 5 × 10^−8^.

## Acknowledgements

D.B. would like to thank Professor Jerzy Ombach for significant help with the process of obtaining access to the data. This research is supported by the European Union's 7th Framework Programme for research, technological development and demonstration under Grant Agreement no 602552, cofinanced by the Polish Ministry of Science and Higher Education under Grant Agreement 2932/7.PR/2013/2 and by NIH grants R01 HG006695, R01MH101782 and R01MH108467.

